# GPCRmd uncovers the dynamics of the 3D-GPCRome

**DOI:** 10.1101/839597

**Authors:** Ismael Rodríguez-Espigares, Mariona Torrens-Fontanals, Johanna K.S. Tiemann, David Aranda-García, Juan Manuel Ramírez-Anguita, Tomasz Maciej Stepniewski, Nathalie Worp, Alejandro Varela-Rial, Adrián Morales-Pastor, Brian Medel Lacruz, Gáspár Pándy-Szekeres, Eduardo Mayol, Toni Giorgino, Jens Carlsson, Xavier Deupi, Slawomir Filipek, Marta Filizola, José Carlos Gómez-Tamayo, Angel Gonzalez, Hugo Gutierrez-de-Teran, Mireia Jimenez, Willem Jespers, Jon Kapla, George Khelashvili, Peter Kolb, Dorota Latek, Maria Marti-Solano, Pierre Matricon, Minos-Timotheos Matsoukas, Przemyslaw Miszta, Mireia Olivella, Laura Perez-Benito, Davide Provasi, Santiago Ríos, Iván Rodríguez-Torrecillas, Jessica Sallander, Agnieszka Sztyler, Nagarajan Vaidehi, Silvana Vasile, Harel Weinstein, Ulrich Zachariae, Peter W. Hildebrand, Gianni De Fabritiis, Ferran Sanz, David E. Gloriam, Arnau Cordomi, Ramon Guixà-González, Jana Selent

## Abstract

G protein-coupled receptors (GPCRs) are involved in numerous physiological processes and are the most frequent targets of approved drugs. The explosion in the number of new 3D molecular structures of GPCRs (3D-GPCRome) during the last decade has greatly advanced the mechanistic understanding and drug design opportunities for this protein family. While experimentally-resolved structures undoubtedly provide valuable snapshots of specific GPCR conformational states, they give only limited information on their flexibility and dynamics associated with function. Molecular dynamics (MD) simulations have become a widely established technique to explore the conformational landscape of proteins at an atomic level. However, the analysis and visualization of MD simulations requires efficient storage resources and specialized software, hence limiting the dissemination of these data to specialists in the field. Here we present the GPCRmd (http://gpcrmd.org/), an online platform that incorporates web-based visualization capabilities as well as a comprehensive and user-friendly analysis toolbox that allows scientists from different disciplines to visualize, analyse and share GPCR MD data. GPCRmd originates from a community-driven effort to create the first open, interactive, and standardized database of GPCR MD simulations. We demonstrate the power of this resource by performing comparative analyses of multiple GPCR simulations on two mechanisms critical to receptor function: internal water networks and sodium ion interaction.

## Introduction

G protein-coupled receptors (GPCRs) are abundant cell surface receptors accounting for ~4% (800) of all human genes. They play a vital role in signal transduction by regulating numerous aspects of human physiology and are the targets of 34% of the drugs approved by the US Food and Drug Administration^1^. Important advances in protein engineering, X-ray crystallography and cryo-electron microscopy (cryo-EM) during the past decade have led to an exponential increase in the number of available GPCR structures (3D-GPCRome) deposited in the Protein Data Bank (PDB) (GPCRdb http://gpcrdb.org/structure/statistics, 2019). This rapid growth has fueled the development of the GPCRdb^2^, an online resource for GPCR reference data, analysis, visualization and data-driven experiment design. This resource provides a wide range of tools including a knowledge-based resource for GPCR crystal and cryo-EM structure determination^3^.

However, static high-resolution structures provide little information on the intrinsic flexibility of GPCRs, a key aspect to fully understand their function. Important advances in the computer science field have transformed computer simulations into a very powerful technique to explore protein conformational landscapes. In particular, all-atom molecular dynamics (MD) simulations have proven useful to complement experiments and characterize GPCR fluctuations at the atomic level^4^. Likely due to technical and sustainability limitations, only a modest number of online resources cover MD simulations (reviewed in 5). Recent large improvements of internet bandwidth, compression of simulation data, and storage capacities now enable faster and larger online repositories that host atom trajectories from MD simulations. Moreover, new visualization^6^ and online file-sharing^7,8^ tools have opened the door to streaming and remotely inspecting MD trajectories online, thereby removing the need for specialized MD software^5^.

Here we present the GPCRmd platform (Fig. 1), the first open-access and community-driven research resource for sharing GPCR MD simulations with the aim of mapping the entire 3D-GPCRome. This new resource paves the way for GPCR scientists from very different disciplines to perform comparative studies on universal aspects of GPCR dynamics. We showcase the potential of GPCRmd for exploring key aspects of GPCR dynamics by performing comparative analyses of internal water molecules and sodium ion binding in multiple GPCR MD simulations. The open and intuitive design of the GPCRmd platform will not only foster interdisciplinary research and data reproducibility, but also transparent and easy dissemination of GPCR MD simulations.

**Fig. 1:**
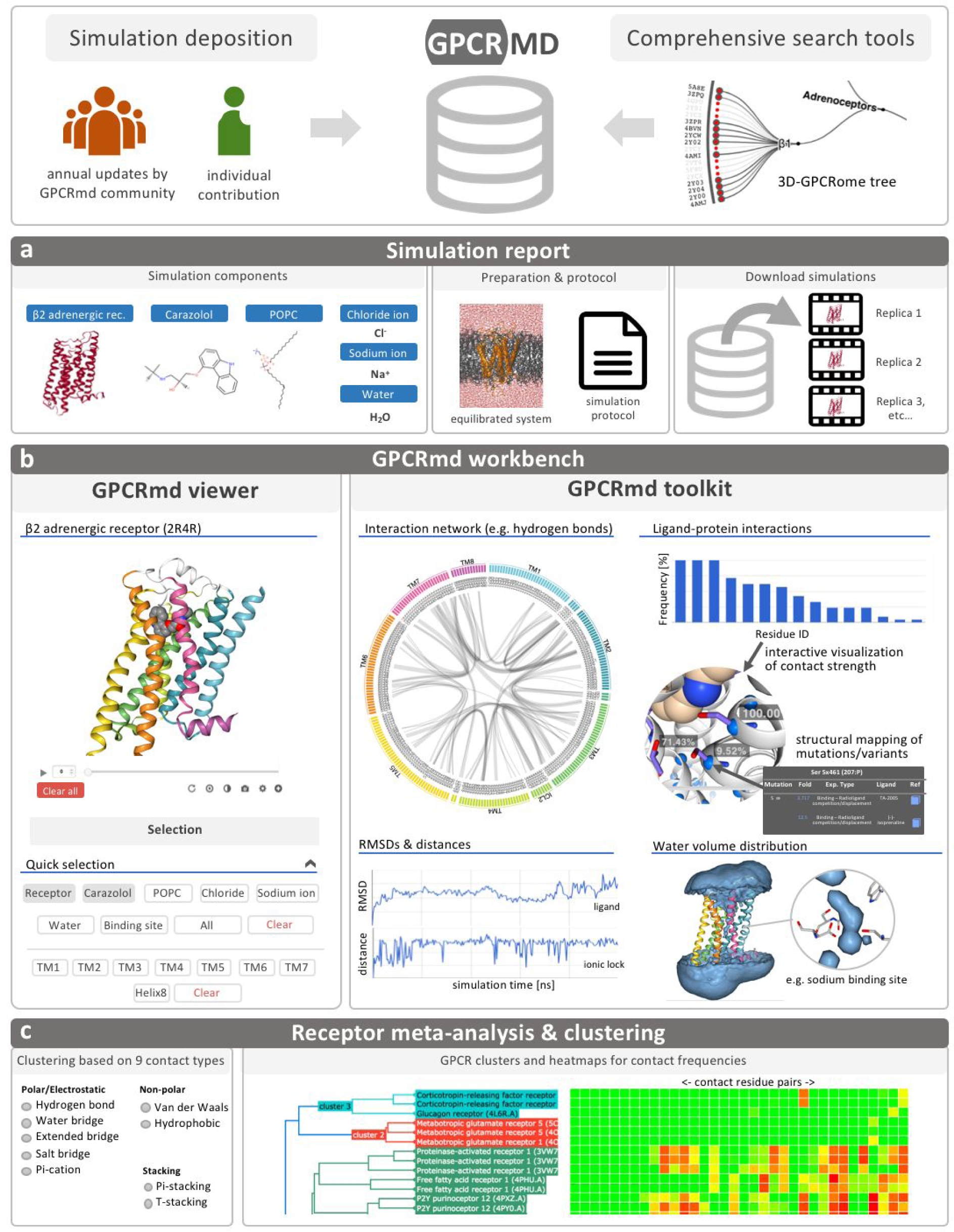
GPCRmd framework. GPCRmd is an online resource for storage, streaming, and analysis of GPCR MD simulation data from individual contributions and annual collective updates. An intuitive search algorithm allows for comprehensive screening of the database. (**a**) The user obtains detailed information about the simulation data via the simulation report. (**b**) A GPCR-specific workbench enables interactive visualization (GPCRmd viewer) and analysis (GPCRmd toolkit) for individual simulations. (**c**) Finally, the comparative analysis and clustering of multiple MD simulations helps finding relationships between receptors based on nine different molecular interaction types.

## Results

### MD simulations from all GPCR classes structurally solved to date

GPCRmd is a community-driven resource that provides direct and interactive visualization of MD trajectories, and that is only contingent on a web browser. As a result, the GPCRmd platform grants easy access for both computational and non-expert scientists. Moreover, we equipped it with a comprehensive set of tools to easily analyse molecular interactions and protein motions involving conserved, pharmacologically relevant, or disease-related residues and structural motifs potentially involved in GPCR function (Fig. 1b,c). In adherence to the Findable, Accessible, Interoperable, and Reusable principles for scientific data management^9^, GPCRmd provides open access to all of its data and simulations protocols (Fig. 1a). Corresponding data are deposited either by individual contributions or annual updates from the GPCRmd community.

We initiated the GPCRmd database by creating a comprehensive MD dataset including at least one representative structure from each of the four structurally characterized GPCR classes. To allow for comparison of ligand-induced effects across receptors, this first set comprises 95 PDB identifiers from 52 different receptor subtypes (Fig. 2) either in their apo form or bound to a natural ligand, surrogate agonist, or antagonist (see Methods). To generate reproducible data, we carefully designed a common protocol for the collective set-up and simulation of all structures listed in Figure 2 (see Methods) and made it publicly available at https://github.com/GPCRmd/MD-protocol. Each system was simulated for 500 ns in three replicates (total time 1.5 µs) allowing for structure relaxation as well as sampling of receptor flexibility. At the time of writing, the GPCRmd platform holds 570 GPCR MD simulations from the GPCRmd community plus 45 simulations from individual contributions totalling to an aggregated simulation time of 408 µs.

**Fig. 2:**
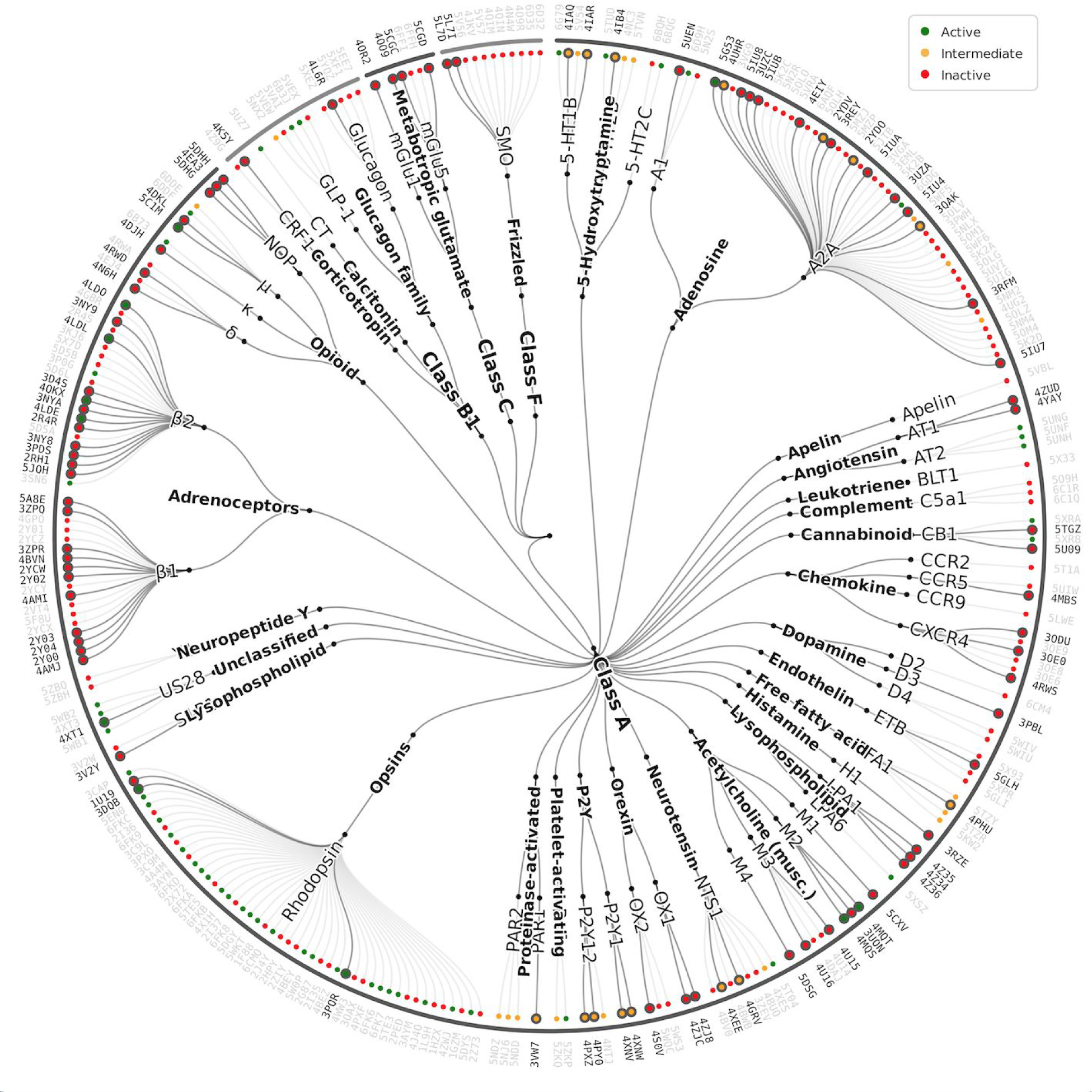
The 3D-GPCRome. Mapping the GPCR structures contained in the first GPCRmd release onto the 3D-GPCRome tree. The first GPCRmd dataset of simulated structures (191 systems at the time of manuscript preparation) covers 100% of GPCR classes, 71% of receptors subtypes, and 80% of GPCR families with solved structure at the time of writing, and accounts for approximately 35% of all GPCR structures deposited in the PDB (black PDB identifiers). Colored circles differentiate between active (green), intermediate (yellow), or inactive (red) receptor states.

### GPCRmd viewer: sharing and interactive visualization of GPCRs in motion

To provide easy sharing and interactive visualization of GPCR MD simulations within the 3D-GPCRome, we created the GPCRmd viewer (Fig. 3). This viewer builds on MDsrv^7^, a recently published tool that allows easy trajectory sharing and makes use of the interactive capabilities of the popular web-based structure viewer NGL^6^.

**Fig. 3:**
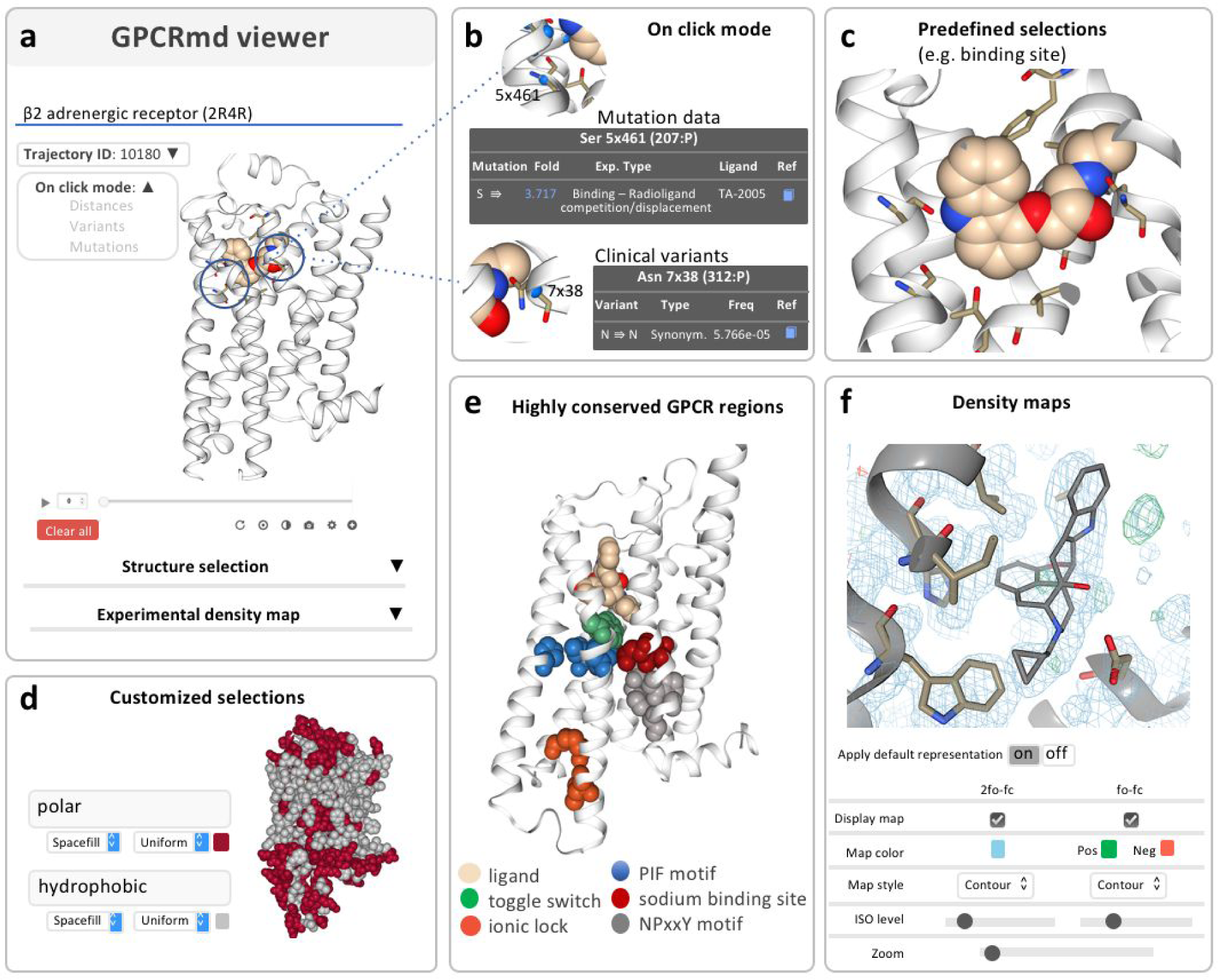
The GPCRmd viewer. Interactive visualization of GPCR MD simulations allows for (**a**) streaming simulations, (**b**) structural mapping of mutation data and clinical variants, (**c**) predefined selections of simulation components and ligand binding sites, (**d**) customized selections that enable tailored visualization of trajectories, (**e**) knowledge-based selections for visualization of GPCR conserved regions, and (**f**) density maps allows for comparison between experiments and MD simulations. A set of predefined, custom and knowledge-based selections enables quick exploration of particular regions of the map such as the ligand binding pocket. Flexible options allow users to change the color of the map type (classical fo-fc or composite 2fo-fc), style (e.g. wireframe / contour), or the surface and zoom levels.

The GPCRmd viewer provides interactive structural analysis of the simulations through on-click actions (Fig. 3b). To account for the fact that almost 25% of the GPCR functional sites show an average of at least one polymorphism, we mapped all GPCR variants^10^ and site-directed mutations^11^ from the GPCRdb^2^ to each GPCR structure. Activation of the modes ‘Show variants’ or ‘Show mutations’ displays, respectively, each variant or mutation as small beads (Fig. 3b). A click on a bead reveals further information on the variant / mutation, including a link to experimental data and the original publication. A separate on-click mode, ‘Show distances’, exploits NGL^6^ to measure atom pair distances.

The powerful selection capabilities of the viewer (Fig. 3c-e) enable fast inspection of trajectories. Standard selections quickly visualize any molecule type in the simulation, neighboring molecules at a custom distance of each other, or specific positions along the protein sequence. It is worth noting that the GPCR viewer makes use of GPCRdb generic residue numbering^12^ by automatically linking each residue to its respective index position. Importantly, predefined knowledge-based selections enable more specific displays such as residues within 2.5 Å of the ligand (Fig. 3c), individual GPCR helices, or highly conserved positions and functional motifs (Fig. 3e). In addition, the NGL selection language (see https://gpcrmd-docs.readthedocs.io/en/latest/) enables the use of custom selection keywords to create tailored representations of any atom or part of the trajectory loaded in the GPCRmd viewer (Fig. 3d). Since several of these keywords stand for the chemical nature or secondary structure of proteins, they are particularly helpful for visual analysis of GPCR dynamics.

Furthermore, the GPCRmd viewer provides visualization of X-ray and electron microscopy (EM) density maps from the PDB. This allows for atomic-level comparison of the GPCR conformational landscape inferred in structurally determined structures and observed in MD simulations (Fig. 3f).

### GPCRmd toolkit: investigation of GPCR dynamics through interactive analysis

The GPCRmd toolkit provides intuitive analysis of the MD simulations by complementing and directly interacting with the GPCRmd viewer (Fig. 1b, left). The toolkit allows to compute custom distances, Root Mean Square Deviation (RMSD), and averaged water density maps for individual simulations (Fig. 1b, right). In addition, it provides interactive tools to qualitatively and quantitatively compare the non-covalent landscape of contacts for the entire GPCRmd dataset (Fig. 1b, right).

#### Interaction network tool

To easily identify relevant non-covalent contacts in GPCRmd simulations, the GPCRmd toolkit uses Flareplots^13^, an interactive circular representation of contact networks that can be displayed per frame or summarized for the complete trajectory (Fig. 4, right). The interaction network tool automatically integrates the GPCRmd viewer with the GPCRmd toolkit, making it straightforward to detect, for instance, differences in the hydrogen bonding network dynamics between active and inactive receptor simulations. The current version of the interaction network tool focuses on intra- and inter-helical interactions including nine different types of non-covalent interactions (see Methods).

**Fig. 4:**
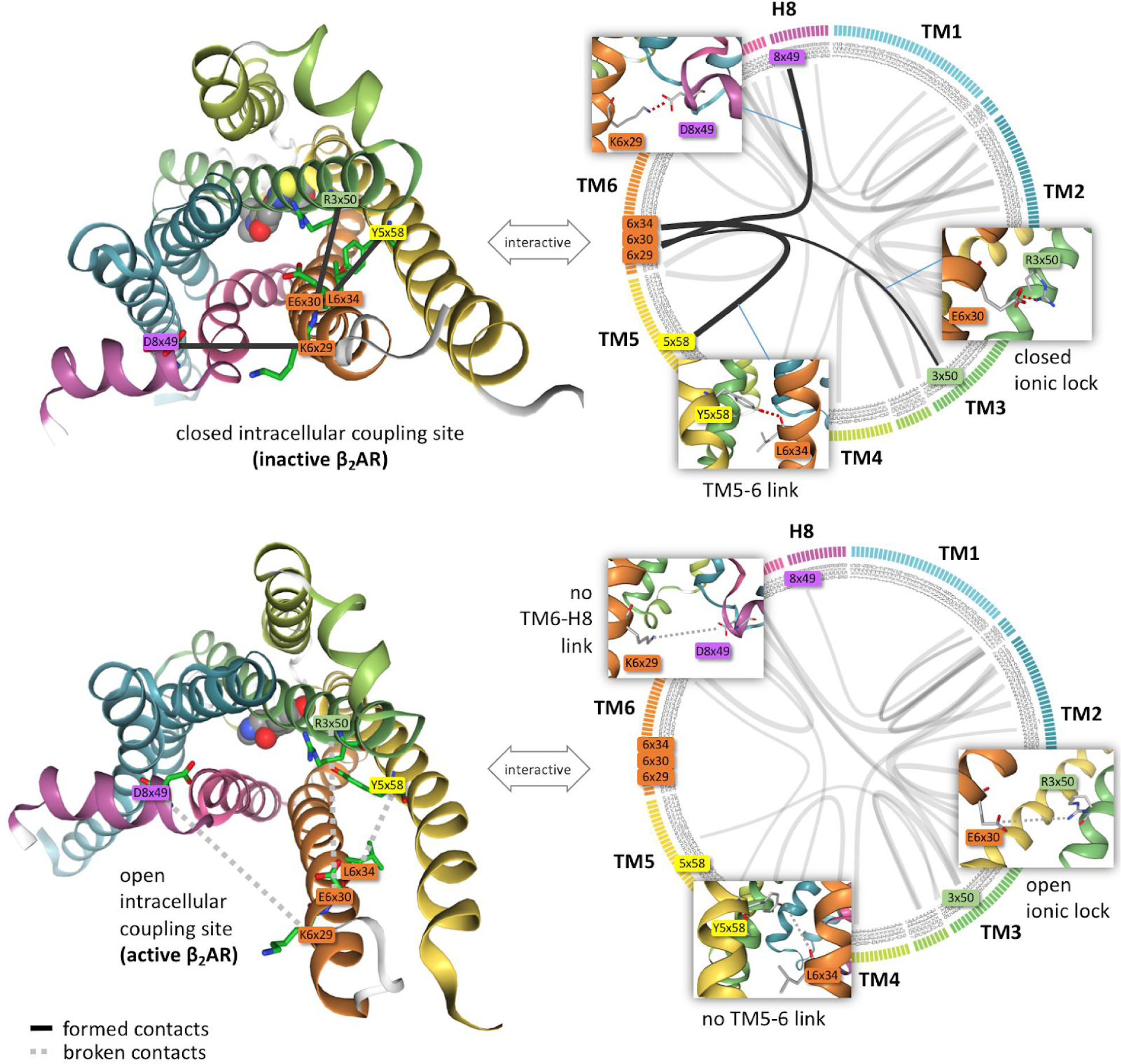
Interaction network tool. Interactive visualization and analysis of intramolecular contacts. Summary plot for the hydrogen bonding network (i.e. average over the entire trajectory) obtained by selecting hydrogen bonds as interaction type. Circular plots (right) for the inactive (upper panel) and active (lower panel) conformations of the β_2_AR, where line thickness represents contact frequency. Comparison of these plots reveals important differences specifically at the intracellular coupling site. The inactive receptor displays contacts that help maintain the receptor in a closed state, such as the characteristic ionic lock between R3×50 and E6×30^14^, a TM5-TM6 linkage established by Y5×58 to the backbone of L6×43, and a TM6-Helix8 connection between K6×29 and D8×49. Such contacts are missing in the active β_2_AR conformation. The user can interactively explore the dynamics of the plotted contacts in the circular plot (right panel) in a structural context (left panel). Residues are numbered according to their GPCRdb generic numbering scheme^12^.

#### Interaction frequency tools

The GPCRmd toolkit provides two dedicated tools to study key electrostatic interactions, namely hydrogen bonds and salt bridges. The hydrogen bonds tool identifies GPCR intra- and intermolecular hydrogen bonds formed during the simulation, whereas the salt bridges tool identifies GPCR intramolecular salt bridges. Moreover, these tools allow studying the interplay between the receptor and the membrane by computing protein-lipid interactions. Furthermore, it can identify protein residues involved in ligand binding through the ligand-receptor contacts tool. The tool outputs the interaction strength at each residue by computing its contact frequency (Fig. 1b, right). All three contact tools provide interactive visualization of the results in the GPCRmd viewer.

#### RMSD tool

The GPCRmd toolkit can monitor a change in distance between any pair of atoms during the simulation. Alternatively, per-frame atom distances can be measured and displayed in the viewer via on-click actions. While distance measurements can provide relevant information on protein structure (e.g. functionally-relevant protein motions, bond formation / breaking, etc.), RMSD calculations are more suited to quantify structural stability and conformational changes. The RMSD tool measures the structural difference of protein and ligand atoms at any point in the simulation with respect to the initial frame. Therefore, it can be used to monitor simulation integrity or structural deviations throughout the simulation. Both tools generate time course plots (Fig. 1b, right) that can interactively link each data point to its respective frame in the viewer.

#### Water volume distribution tool

Due to the vital role of internal water molecules in GPCRs^15^, we equipped the GPCRmd toolkit with a water density map tool. This tool can quickly display an averaged water density map of the MD trajectory under study in the GPCRmd viewer (Fig. 1b, right), thus allowing to monitor, for example, the formation of the continuous internal water channel known to be essential for GPCR activation^16^.

### Functional hotspots discovered through meta-analysis of GPCR simulations

The GPCRmd platform can uniquely compare GPCR simulations within the 3D-GPCRome (Figs. 1c and 2). We developed a module specifically comparing multifold GPCR simulations to uncover universal or distinct mechanisms governing the structural dynamics of these receptors. This module computes the contact frequency of each residue pair for multiple simulations and displays a global comparative analysis via an interactive heatmap plot (Fig. 5a, left). The tool also performs clustering analysis of the contact frequency data to hierarchically classify each receptor and display the resulting tree alongside the heatmap plot (Fig. 5a, left). To further facilitate the interpretation of large heatmaps, we added interactive analysis and visualization capabilities of selected clusters using Flareplots^13^ and NGL^6^.

**Fig. 5:**
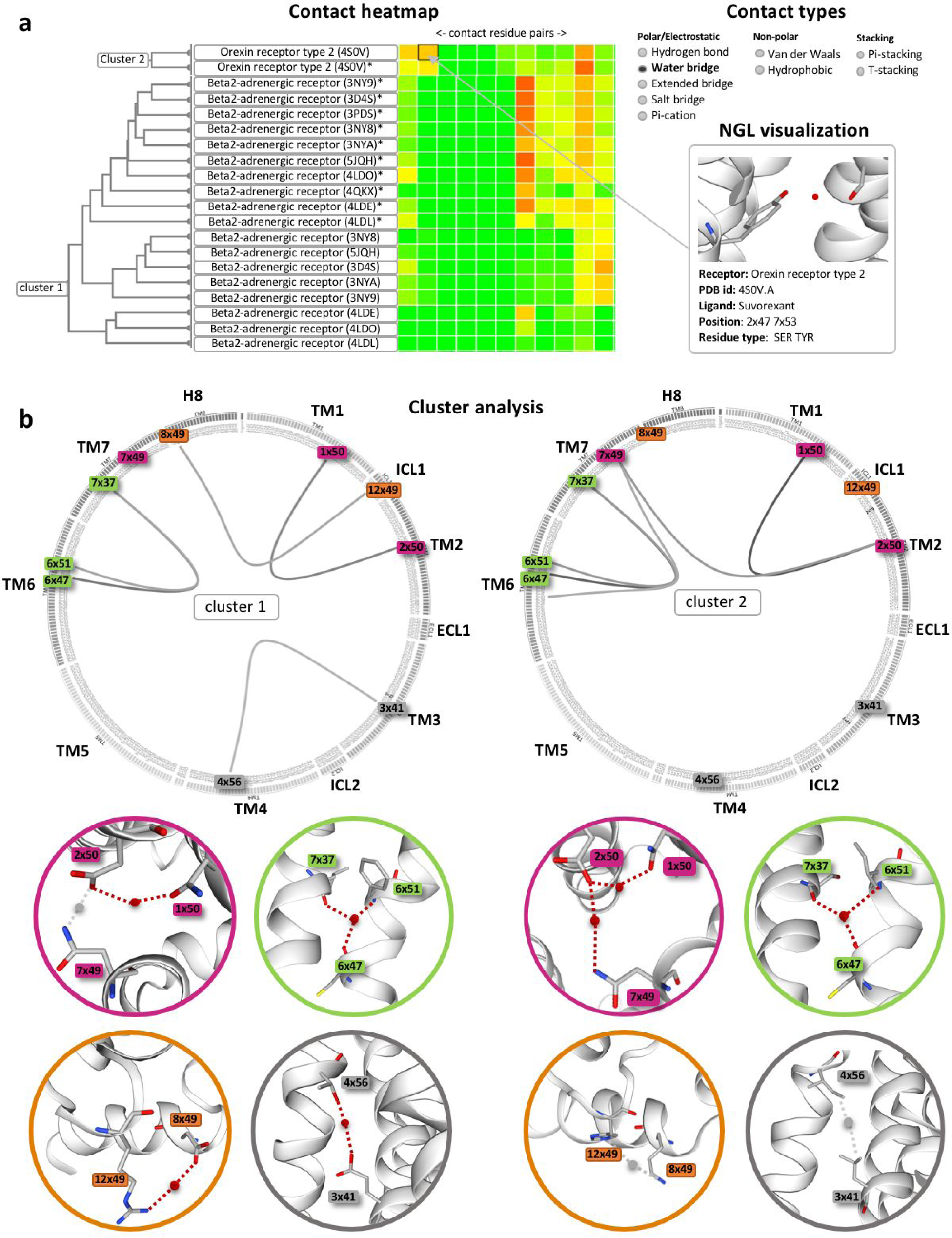
A water bridge signature revealed by comparative analysis using the GPCRmd. (**a**) Heatmap of water-mediated interactions of clusters belonging to the β2AR and OX2R. The plot displays each residue pair (columns) for each GPCR (rows). Green to red color scale stands for low to high contact frequency. Users can select up to nine different non-covalent interaction types to perform the analysis across the complete GPCRmd database or just using a custom subset of simulations. On-click actions provide detailed information on the specific interaction and system involved for each cell of the heatmap. (**b**) Representative water-mediated interactions for the investigated clusters are shown in circular plots. Corresponding structural depictions of interactions are found below the circular plots. This includes a water-mediated network connecting the allosteric sodium binding site 2×50 in TM2 with position 1×50 in TM1 in the β2AR and OX2R (highlighted in purple). This water network is extended from TM2 (2×50) to TM7 (7×49) in the OX2R cluster. Such a water network extension is not observed in the β2AR cluster due to closer proximity of residues 2×50 and 7×49, which enables direct, unmediated, contacts. Another conserved water-mediated feature is a bifurcated polar network linking TM6 (6×47, 6×51) and TM7 (7×37) via helix backbones in the β2AR and the OX2R clusters (highlighted in green). Important differences between both clusters are two water-mediated connections, namely ICL1 (12×49) - H8 (8×49) (highlighted in orange) and TM3 (3×41) - TM4 (4×56) (highlighted in grey), the latter one occurring in a region of the receptor facing the membrane and exclusively found in the β2AR cluster. Residues are numbered according to their GPCRdb generic numbering scheme^12^.

To demonstrate the utility of the meta-analysis tool, and due to their critical role in receptor function^17,18,15,16^, we investigated the interaction fingerprint of water molecules in GPCRs. Along with previously described^19^ conserved water networks, this analysis revealed other water-mediated interactions that are conserved among different receptor subtypes and firstly reported here. For example, in line with Venkatakrishnan et al.^19^, the β2-adrenoceptor (β2AR) and OX_2_-receptor (OX2R) display a common water network that links TM1 (N1×50) and TM2 (D2×50) and a bifurcated network connecting TM6 (6×47, 6×51) and TM7 (7×37) (Fig. 5). Our study shows that this bifurcated network is less prominent in active structures (Supplementary Figure 8). Taking into account that TM6 undergoes large conformational changes upon receptor activation, it is tempting to speculate that uncoupling the interactions between individual water molecules in this bifurcated network represents a step during receptor activation.

Likewise, our analysis reveals a water bridge between intracellular loop 1 (ICL1) and helix 8 (H8) only present in the β2AR (Fig. 5b, right). Further studies (e.g. site-directed mutagenesis) could be used in the future to investigate whether this water bridge contributes to the distinct coupling efficacy and/or specificity shown by the β2AR (principal signaling pathway: Gs family^20^) and OX2R (principal signaling pathway: Gs family, Gi/Go family, Gq/G11 family^20^). Finally, our collective analysis reveals a water bridge between TM3 (3×41) and TM4 (4×56) only found in the β2AR (Fig. 5b, right) and likely related to the striking change in receptor stability observed upon mutation of residue E3×41 in experiments^21^.

### Exploiting the entire GPCRmd dataset: custom analysis of sodium ion interactions across class A GPCRs

We made the entire GPCRmd dataset available for download (see Methods), thus opening the door for the scientific community to perform comparative analyses of multiple simulations across several receptor structures, families, subtypes and classes. To demonstrate the value of such a comprehensive dataset, we studied sodium ion (Na^+^) interaction in GPCRs^22^, an almost universal, albeit poorly understood, mechanism of allosteric modulation of this receptors^23^. We analysed Na+ interaction to conserved orthosteric (3×32) and allosteric (2×50) residues in 261 simulations (87 different apo structures x 3 replicas) covering 47 different class A receptor subtypes. The markedly different frequencies of Na^+^ interaction with these two residues enable receptors to be clustered in three groups (I, II, and III, Fig. 6a,c). Note that our dataset (3 x 0.5 μs) provides valuable insights into sodium interaction sites but it is not sufficient to conclude about binding kinetics.

**Fig. 6:**
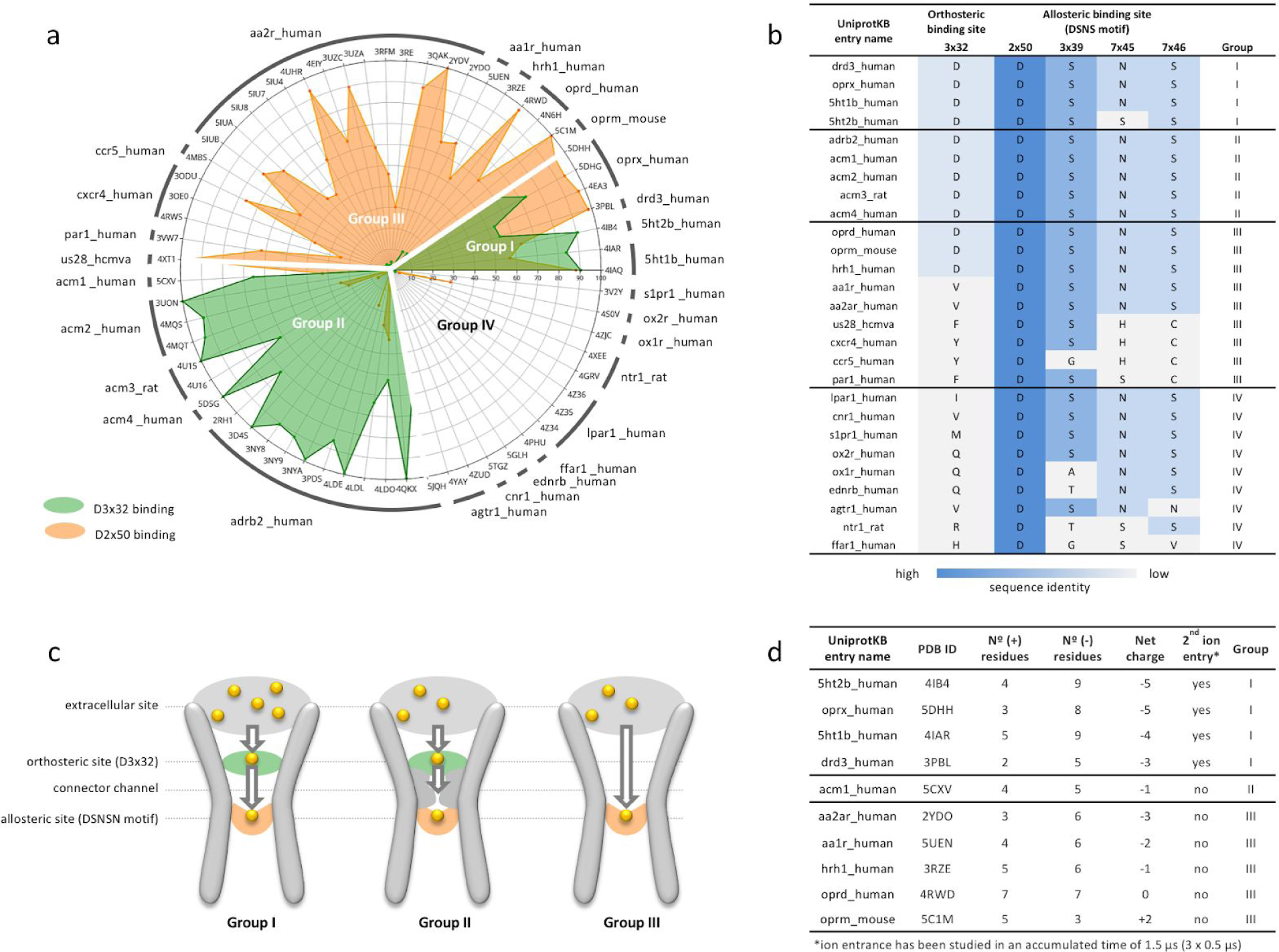
Allosteric Na+ ion interaction in class A GPCRs. (**a**) Na+ interaction frequency at D3×32 (green) and D2×50 (orange) in class A GPCRs across 63 structures including 27 different receptor subtypes. Receptor subtypes and 3D structures are identified by UniprotKB and PDB identifiers, respectively. The radar plot shows the prevalence of sodium interactions (0 to 100%) over the total accumulated simulation time of 1.5 µs (3 x 0.5 µs). (**b**) Sequence alignment of sodium binding sites for the GPCR subtypes included in the simulated dataset. Allosteric binding consists of a multi-step binding process typically initiated with accumulation at the extracellular receptor side followed by receptor penetration through the orthosteric binding site (visiting D3×32, if present) before progressing to the allosteric site D2×50. (**c**) GPCRs can be classified into three groups based on the sodium interaction profile. The interaction profile is driven by the structural features of the sodium entrance channel. (**d**) Extracellular net charge and receptor entrance of a second ion.

In line with previous studies using multiple simulations^24^, our analysis shows that Na+ binds to D2×50 and/or position 3×32 in most of the receptor subtypes (Fig. 6a). Group I (serotonin, dopamine and nociception receptors) shows high sodium interaction frequencies to positions 3×32 and 2×50 and, the latter being stabilized by a hydrogen bonding network often composed of D2×50, S3×39, N7×45 and S7×46 (so-called as DSNS motif) (Fig. 6b). The high interaction frequency to both positions implicates that at times Na+ ions bind simultaneously to position 3×32 and the allosteric site at D2×50. This seems to be a consequence of a higher negative net charge at the extracellular side (Fig. 3c,d), which increases the local concentration of positively charged Na+ around the receptor entrance, and likely facilitates the simultaneous entrance of a second ion. Notably, despite a completely conserved DSNS motif, group II (β-adrenergic and muscarinic receptors) shows surprisingly marginal interaction frequency at D2×50, while still exhibiting a high interaction frequency at 3×32. Visual inspection of the simulation reveals hydrophobic barriers that hamper Na+ passage from 3×32 to 2×50 (Fig. 6c), in line with previous MD simulation studies^24,25^. In contrast to group II, we find high interaction frequencies at position 2×50 for group III and none or only marginal contacts with 3×32. Most receptors of this group (e.g. adenosine A_1A_ and A_2A_, or chemokine CCR5 and CXCR4 receptors) lack an aspartate in position 3×32 allowing for direct diffusion to position 2×50 (Fig. 6c). Interestingly, despite having a D3×32 (Fig. 6b), only low binding is observed during the simulated time frame at this position for a small subgroup of receptors including histamine H_1_ and opioid μ and δ receptors. More simulation time would be required to improve the sampling of ion binding to D3×32. Finally, in a particular subset of receptors, Na+ binds neither to D2×50 nor to position 3×32 within the studied time frame (group IV, Fig. 6a). In fact, slower Na+ binding kinetics has previously been reported^24^ and could be the consequence of blocked access to the binding site from the extracellular side (e.g. receptors taking up ligands from the lipid bilayer).

While our results confirm the essential role of D2×50 for allosteric sodium binding^15,26^ in class A GPCRs, they also reveal that the presence or absence of D3×32 in the orthosteric binding site determine distinct Na+ binding profiles. This analysis exemplifies the potential of the comprehensive GPCRmd dataset to investigate how GPCR sequence, structure and dynamics can jointly contribute to receptor allosteric modulation.

## Discussion

In the last decade, static structures in the 3D-GPCRome have predominantly been described as either active, intermediate, or inactive states. However, a growing body of research suggests that GPCRs are not two- or three-state systems but exhibit a wide range of conformational states with sometimes subtle yet important differences. While several experimental techniques such as nuclear magnetic resonance (NMR)^27^, double electron-electron resonance (DEER)^28^, or single-molecule fluorescence energy transfer (smFRET)^29^ have provided relevant insights into the dynamics and flexibility of GPCRs, MD simulations have emerged as the most promising opportunity to study the complexity of GPCR conformational dynamics in atomistic detail^4^. Moreover, MD simulations can resolve mechanistic elements at time scales and conditions that are not always accessible with experimental techniques.

We have demonstrated the utility of the GPCRmd platform by performing comparative analyses across multiple receptors of two relevant aspects of GPCR biology, namely water network and allosteric Na+ interaction analysis. Using GPCRmd tools, we pinpointed differences in the water-mediated networks of the OX2R and the β2AR potentially involved in receptor activation and G protein coupling. In addition, we showcase the power of exploiting the GPCRmd dataset offline by downloading a comprehensive group of class A GPCR simulations and using external means to investigate Na+ interactions. This study allowed to classify receptors in different groups based on the interaction profile of Na+ to the orthosteric and/or allosteric sites. Interestingly, our study suggests that the probability of ion entrance into the orthosteric site is modulated by the extracellular net charge of the receptor.

### A platform for interdisciplinary investigation of the 3D-GPCRome

The GPCRmd is designed to facilitate interactions and data exchange between GPCR scientists of different disciplines including structural and evolutionary biologists, computational and medicinal chemists, protein engineers, and structural biologists (Table 1). Our tool will become a useful asset for experimental laboratories by providing open access to the dynamic context of specific GPCRs, hence directing or assisting functionally relevant experiments such as cross-linking or mutagenesis studies. Similarly, protein engineers and structural biologists will now be able to employ the GPCRmd workbench to quickly identify specific flexible regions that potentially require protein stabilization.

**Table 1.** Examples of how researchers from different scientific disciplines can make use of the GPCRmd database.

| User | Usage | GPCRmd features | Added value |
| --- | --- | --- | --- |
| **Protein engineers** | **Stabilizing proteins for crystallization.**<br>Detection of flexible receptor regions that require stabilization to improve crystallization success. | GPCRmd workbench including the GPCRmd viewer and Toolkit | Flexible receptor regions are poorly captured in experimental density maps |
| **Crystallographers** | **Retrospective refinement of experimental density maps**<br>(i) Detection of highly flexible regions that explain low-resolution regions in experimental density maps.<br>(ii) Detection of stable water or ion binding sites that can explain unmatched electron density areas.<br>(iii) Rotamers and protonation states. | GPCRmd viewer. MD streaming with overlaid experimental density maps<br>The water map tool in the GPCRmd toolkit | Water and ion binding are poorly captured in experimental density maps<br>Rotation states and corresponding protonation states are difficult to obtain from experimental density maps |
| **Biophysicists** | **Study of interaction networks critical for receptor functionality.** Search for receptor regions to implement linkers or signaling probes (FRET, NMR, etc.) to study receptor functionality. | GPCRmd workbench including the GPCRmd viewer and Toolkit | Receptor dynamics is not available in experimental density maps |
| **Evolutionary biologists** | **Structural relationships and diversity across different GPCRs.** How does evolution (i.e. small sequence differences) impact receptor dynamics. | Comparative receptor analysis and clustering tool | Receptor dynamics is not available in experimental density maps |
| **Medicinal chemists & drug designers** | **Improvement of drug-receptor interactions and design of new drugs.**<br>(i) Exploration of ligand-receptor contacts.<br>(ii) Detection of indirect interactions (i.e. water or ion-mediated) that are crucial for ligand binding.<br>(iii) Flexible and transient switches | Ligand-receptor contacts and water volume distribution tools | Stability of ligand-receptor interactions cannot be deduced from experimental density maps |
| **Biomedical researcher and clinicians for personalized medicine** | **Design of treatment strategies for personalized medicine.** Estimation of the impact of polymorphisms / variants on drug response through their ability to alter drug-receptor interactions or receptor dynamics in regions relevant for receptor functionality (e.g. PIF motif, G protein coupling site). | Cross-linked mutation and variant information | The impact of polymorphism/variants on the strength of ligand-receptor contacts or receptor dynamics cannot be deduced from static structures |
| **Computational biologists (MD novices and experts, bioinformaticians)** | (i) Aid in experimental design and comparison of setup and results in terms of force field performance, impact of ligands, or mutations.<br>(ii) Support for modelling dynamic regions able to adopt distinct conformations.<br>(iii) Guide docking experiments based on the sampled conformational space. | GPCRmd viewer for simulation streaming<br>Simulation protocol and input structures<br>GPCRmd workbench including the GPCRmd viewer and Toolkit | Receptor dynamics are not available in experimental density maps |
| **Students & teachers** | **Visually learning about protein dynamics:** e.g. receptor inactivation, water channel formation in active receptor structures, allosteric binding of sodium ions. | GPCRmd workbench including the GPCRmd viewer and Toolkit | Dynamics cannot be visualized in printed form and trajectories are not part of additional teaching materials |
| **Reviewers and publishers** | Data made available for scrutiny of molecular dynamics articles | GPCRmd platform | Transparency and reproducibility |

Moreover, the GPCRmd will be of great benefit to medicinal chemists and drug designers. For the first time, they will be able to quickly use atomic-level information on the stability / strength of specific ligand-receptor interactions, and the binding of water molecules or ions using the ligand-receptor contacts or water volume distribution tools (see Fig. 1 and Table 1). In addition, drug design scientists can use GPCRmd to investigate potential ligand binding and unbinding pathways based on the dynamics of specific structural elements such as loops, hence aiding the design of new or improved compounds. Furthermore, the GPCRmd can provide valuable structural insights into the location of natural variants and its potential impact on drug binding or receptor functionality. Our cross-referenced data allows easy mapping of variants and site-directed mutations onto the receptor structure and investigation of their dynamics during the simulations (Fig. 1b, right, and Fig. 4b). This could guide further investigations to predict drug efficacy or adverse reactions in individuals with a specific variant and in turn support the selection of more efficacious and safer drug treatments.

Beyond wet-lab applications, GPCRmd is an important dissemination resource for computational biologists, ranging from students and MD novices to MD experts and bioinformaticians from related fields. Our platform offers a harmonized database to perform future comparative studies across different MD setups, force fields, ligands, lipid compositions or GPCR variants, which offers a significant advantage over currently available archives or data repositories such as FigShare (https://figshare.com/) or Zenodo (http://zenodo.org).

### The GPCRmd consortium: reproducible and sustainable research in GPCR MD simulations

This community-driven effort has laid the foundation of the GPCRmd consortium, an open community of GPCR computational researchers driving the centralization, dissemination, and development of open-source and reproducible analysis of massive amounts of GPCR MD data. We believe that GPCRmd will enhance the dissemination of scientific results by offering a platform to make published protocols and simulation data publicly available. This will promote transparency, consistency, and reproducibility in the field of GPCR dynamics. On the other hand, community engagement will overcome one of the most important challenges faced by this kind of resources, namely sustainability. The implementation of the GPCRmd consortium under the umbrella of the active European Research Network on Signal Transduction (ERNEST, https://ernest-gpcr.eu) will provide support to increase the coverage of the 3D-GPCRome with future releases of the GPCRmd platform. While the first GPCRmd dataset from the consortium already maps more than 70% of GPCR subtypes within the 3D-GPCRome, future annual releases will further increase this coverage bridging the gap between solved and simulated structures. Furthermore, individual contributions will provide extra valuable insights into specific key features of GPCRs such as their ion permeation capacity^30^, their ability to act as constitutive phospholipid scramblases^31^ or ligand-dependent receptor activation pathway^32^.

## Methods

### MD simulations

The first GPCRmd includes 95 different GPCR structures either bound to their natural ligand (e.g. sphingosine-bound S1P_1_R), an agonist (e.g. ergotamine-bound 5HT_2B_R), or an antagonist (e.g. alprenolol-bound β_2_AR). In addition to ligand-bound structures, we included an apo form of each receptor by removing the ligand from its binding pocket. We carefully designed a common protocol for the collective set-up and simulation (Supplementary Note A) phases of all structures. During the set-up phase, different expert members of the GPCR-MD community individually prepared each family of GPCR structures by refining / remodeling Protein Data Bank (PDB) structures (e.g. missing residues, disulfide bridges, co-crystallization molecules, loop remodeling, etc.), placing missing water molecules and sodium ions, or assigning relevant protonation states (Supplementary Note A). Next, each protein was prepared for simulation by embedding it in a lipid bilayer and adding water and ions to the system. Each system was equilibrated following a standard procedure previously outlined and discussed within the GPCR-MD community (Supplementary Note A). Finally, the distributed computing platform GPUGRID^33^ was used to simulate 3 replicas of each system for 500 ns (i.e. accumulated 1.5 μs). We made all set-up and simulation protocols openly available at https://github.com/GPCRmd/MD-protocol.

### Database structure

The GPCRmd database and web interface were developed using Django Web Framework (v1.9), Python (v3.4), JavaScript libraries, jQuery 1.9, jQuery UI 1.11.2, and PostgreSQL 9. The structure of the database is based on five main objects: protein objects identified by their sequence and their relationship with UniprotKB entries (Supplementary Fig. 1), molecular entities (molecule object) identified by an InChI^34^ generated with forced hydrogen connectivity (Supplementary Fig. 2), crystalized assembly (model) (Supplementary Fig. 3), molecular dynamics simulations (dynamics) objects (Supplementary Fig. 4), and chemical species (compound) identified by standard InChI. Supplementary Fig. 5 shows the Entity Relationship (ER) diagram. Furthermore, we incorporated experimental data from IUPHAR^35^ and BindingDB^36^, and linked each main object to bibliographic references. GPCRdb^2^ tables were used to add standard nomenclatures to GPCR sequence residue numbers.

### Custom analysis

The whole GPCRmd repository is released as open source under the https://creativecommons.org/licenses/by/4.0/ hence enabling downloading and custom analysis of the comprehensive dataset. Each trajectory can be downloaded from its respective link at the simulation report page (see https://gpcrmd-docs.readthedocs.io/en/latest/). We exemplified this usage by studying sodium ion binding across a selection of class A GPCRs within the GPCRmd dataset. The frequency of sodium ion binding to the closest oxygen atom of the carboxylic group (2 x O^ϵ^) of residues 3×32 and 2×50 were computed using a cutoff distance of 5 Å. Both highly conserved positions are normally aspartates. For non-conserved residues we used the following atoms: Gln (N^ϵ^, O^ϵ^), His (N^δ^), Arg (N^ϵ^), Ala (C^γ^), Val (2 x C^γ^ Hydrogen), Ile (2 x C^γ^ Hydrogen), Met (S^δ^), Phe (2 x C^δ^), Tyr (2 x C^δ^).

### GPCRmd viewer

The GPCR viewer uses builds on NGL^6^ and MDsrv^7^ and uses data from the PDB (rcsb.org^37^), the GPCRdb^2^, and the ExAC database^38^.

#### On-click modes

Data for on-click variants and site-directed mutagenesis annotations are taken from the GPCRdb^2,10,11^ and include: generic GPCR numbers^12^, original and mutated residues, effect of the mutation in ligand binding (fold change), experiment type, ligand used for the experiment, and bibliographic reference. Variant data is obtained from the ExAC database^38^, and includes amino acid substitutions (canonical and variant), allele frequencies, and link to the ExAC entry describing the variant. On-click selection capabilities build on NGL^6^ web viewer, which allow the creation of different representation objects using the http://nglviewer.org/ngl/api/manual/selection-language.html. GPCRmd selection capabilities also feature the GPCR generic numbering scheme^12^. In this case, GPCRdb numbers are adapted to the http://nglviewer.org/ngl/api/manual/selection-language.html through regular expressions.

*Experimental density maps are* loaded from PDB and aligned to the first frame of the simulation displayed using NGL^6^. The transformation matrix applied to the density map in order to perform the alignment is pre-computed using the Python library MDAnalysis^39^.

### GPCRmd toolkit

#### Interaction networks

Non-covalent residue-residue interactions formed in the simulation are displayed using *Flareplots*^13^. To pre-compute interactions during the simulation, we used *GetContacts*^13^ in all interaction types except for hydrogen bonds, where we used the definition of Wernet and Nilsson. We manually integrated Flareplots and NGL to allow for interactivity between the GPCRmd toolkit and the GPCRmd viewer.

#### Interaction frequencies

*Hydrogen bonds* are calculated using the “*wernet_nilsson*” module of MDtraj^40^. A hydrogen bond is defined using distance and angle cut offs between hydrogen donor (NH or OH) and acceptor (N or O) atoms as follows:

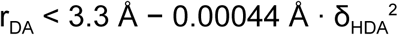

where r_DA_ is the distance (Å) between donor and acceptor heavy atoms, and δ_HDA_ is the angle (degrees) formed between the hydrogen atoms of donor and acceptor atoms. By default, the analysis does not consider hydrogen bonds between neighbouring residues and includes side chains as well as backbone atoms. *Ligand-receptor contacts* are computed using the *compute_contacts* module of MDtraj^40^. *Salt bridge frequency* is computed using the “*compute_distances*” module of MDtraj^40^. Salt bridges are defined as any combination between the sets {Arg-NH1, Arg-NH2, Lys-NZ, His-NE2, His-ND1} and {Glu-OE1, Glu-OE2, Asp-OD1, Asp-OD2} with atoms closer than 4 Å. Histidine atoms are only considered if the residue is protonated. The *distance* between atom pairs through the entire or strided trajectories is computed using the “*compute_distances"* module of MDtraj^40^. Atom pairs can be defined either using the “Show distances” on-click mode and imported to the tool, or http://nglviewer.org/ngl/api/manual/selection-language.html instances.

<u>*RMSD*</u> is computed using the *rmsd* module of MDtraj^40^. The first frame of the trajectory is used as a reference structure by default. The atoms used for RMSD computation can be defined using the provided pre-selection in the GPCRmd toolkit (e.g. protein alpha carbons, non-hydrogen protein atoms, ligand, etc.). RMSD is computed after optimal alignment according to the following equation:

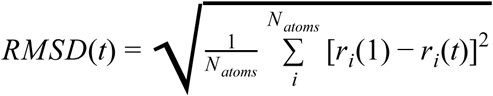

where *N*_*atoms*_ is the number of atoms for structure comparison, *r*_*i*_*(1)* is the position of atom *i* in the reference frame (i.e. trajectory frame 1) and *r*_*i*_*(t)* is the position of atom *i* at time *t* of the trajectory.

#### Water volume distribution

Water occupancy maps are pre-computed and stored on the server side using the VolMap tool of VMD^41^. Maps are generated only for oxygen atoms of a water molecule using a cutoff distance of 10 Å to the protein and a resolution of 1 Å. Atoms are treated as spheres using their atomic radii. The resulting isosurface is displayed in the GPCRmd viewer.

### Meta-analysis tool

Contacts are computed using *GetContacts*^13^ and results plotted as interactive heatmaps using the Bokeh visualization library (https://docs.bokeh.org/en/latest/). Contact frequencies per system are averaged over simulation replicas. For accurate comparison, residue contact pairs are aligned using the GPCRdb generic numbering scheme^12^. Hierarchical clustering uses the “linkage” function of the scipy^42^ library with default parameters. Dendrogram plots use the Plotly library (https://plot.ly/python/).

## Acknowledgements

The GPCRmd consortium acknowledges the support of COST Action CA18133, the European Research Network on Signal Transduction (https://ernest-gpcr.eu) and COST Action CM1207 GLISTEN. The authors would like to thank Rasmus Fonseca and AJ Venkatakrishnan for their help implement Flareplots into the GPCRmd toolkit. MTF acknowledges financial support from the Spanish Ministry of Science, Innovation and Universities (FPU16/01209). TMS would like to acknowledge support from the National Center of Science, Poland (grant number 2017/27/N/NZ2/02571). IRE acknowledges Secretaria d’Universitats i Recerca del Departament d’Economia i Coneixement de la Generalitat de Catalunya (2015 FI_B00145) for its financial support. PK thanks the German Research Foundation DFG for Heisenberg professorship KO4095/4-1 and KO4095/5-1 as well as project KO4095/3-1 (funding MMS). GDF acknowledges support from MINECO (Unidad de Excelencia María de Maeztu MDM-2014-0370 and BIO2017-82628-P) and FEDER and from the European Union’s Horizon 2020 Research and Innovation Programme under Grant Agreement No. 823712 (CompBioMed2 Project). DL would like to acknowledge support from the National Centre of Science in Poland (DEC-2012/07/D/NZ1/04244). PWH thanks the DFG (Hi 1502, SFB 1423 / Z04), the Stiftung Charité and the Einstein Foundation. SF thanks the National Science Centre Poland grant no 2017/25/B/NZ7/02788. JKST likes to acknowledge support from HPC-EUROPA3 (INFRAIA-2016-1-730897) and the EC Research Innovation Action under the H2020 Programme. The work was supported by grants from the Swedish Research Council (2017-4676), the Swedish strategic research program eSSENCE, and the Science for Life Laboratory to J.C. HW and GK acknowledge support from NSF grant #1740990 for In Situ Data Analytics for Next Generation Molecular Dynamics Workflows, and the 1923 Fund. Finally, JS acknowledges financial support from the Instituto de Salud Carlos III FEDER (PI15/00460 and PI18/00094) and the ERA-NET NEURON & Ministry of Economy, Industry and Competitiveness (AC18/00030).

## Author contributions

Conceptualization: JS, RGG, IRE, MTF; Database structure: IRE; GPCRmd workbench: MTF with support from NW, AVR and FS; Meta-analysis tool: DAG with support from MTF and IRE; Submission system: JMAR with support from IRE; Query system: AVR with support from IRE; Server maintenance: MTF; Simulation standard protocol - original draft: RGG and JS; Simulation standard protocol - revision: GDF, AC, IRE, JC, HDT, JW, MMS, PK, JKST, PWH, TMS, SF, TG, MJR; Protein curation - modelling missing loops: GPS and DEG; Protein curation - placement of internal water molecules: EM, PWH and AC; Protein curation - expert knowledge for final curation (e.g. protonation states, disulfide bridges, etc.): IRE, MTF, JKST, DAG, JMRA, TMS, NW, AVR, AMP, BML, GPS, EM, TG, JC, XD, SF, JCGT, AG, HGDT, MJR, WJ, JK, PK, DL, MMS, PM, MTM, PM, MO, LPB, SR, IRT, JS, AS, SV, PWH, GDF, FS, DEG, AC, RGG, JS; Coordination of data exchange: TMS; Preparation of solvated receptor-membrane systems: IRE with support from TMS; Molecular dynamics simulation: GDF, IRE and BML; MD data curation and submission: IRE, AMP, MTF, DAG, TMS, and JS; Individual contribution of MD data: GK, HW, UZ, NV, DP and MF; Manuscript writing - Original Draft: RGG, JS with input from IRE, MTF and JKST; Manuscript - Review & Editing: all authors with important contributions from DEG, TG, and PK; Project supervision and administration: JS.

