## Supplementary Information for "GPCRmd uncovers the dynamics of the 3D-GPCRome"

[9]*Department of Biosciences, University of Milan, Milan, Italy*
[10]*Science for Life Laboratory, Department of Cell and Molecular Biology, Uppsala University, Uppsala, Sweden*
[11]*Laboratory of Biomolecular Research, Paul Scherrer Institute (PSI), 5232 Villigen PSI, Switzerland.*
[12]*Condensed Matter Theory Group, Paul Scherrer Institute (PSI), 5232 Villigen PSI, Switzerland.*
[13]*Faculty of Chemistry, Biological and Chemical Research Centre, University of Warsaw, Warsaw, Poland*
[14]*Department of Pharmacological Sciences, Icahn School of Medicine at Mount Sinai, New York, NY, USA*
[15]*Department of Cell and Molecular Biology, Uppsala University, Biomedical Center, Box 596, SE-751 24, Uppsala, Sweden*
[16]*Department of Physiology and Biophysics, Weill Cornell Medical College of Cornell University, New York, NY 10065, USA*
[17]*Institute for Computational Biomedicine, Weill Cornell Medical College of Cornell University, New York, NY 10065, USA*
[18]*Department of Pharmaceutical Chemistry, Philipps-University Marburg, Marbacher Weg 6, 35032 Marburg, Germany*
[21]*MRC Laboratory of Molecular Biology, Francis Crick Avenue, Cambridge CB2 0QH, UK*
[22]*Department of Pharmacy, University of Patras, 26504, Patras, Greece*
[23]*Department of Computational and Quantitative Medicine, Beckman Research Institute of the City of Hope, Duarte, California*
[24]*Computational Biology, School of Life Sciences, University of Dundee, Dundee, UK*
[25]*Physics, School of Science and Engineering, University of Dundee, Dundee, UK*
[26]*Berlin Institute of Health, 10178 Berlin, Germany*
[*] These authors contributed equally.

**Supplementary Note A. System set-up and simulation protocol**

*Protein structure preparation*. The structures of simulated GPCRs were obtained from the Protein Data Bank (PDB) (rcsb.org[1]) (see Table S1 for PDB IDs). Auxiliary proteins and long unresolved or truncated N- and C-terminal regions were removed and stabilizing mutations were reverted back to the native sequence. In receptors where ICL3 was longer than 10 residues, a chain break was introduced in the middle and five residues at each end were modeled using MODELLER[2]. Modelling and refinement of the structures was performed using the methodology described by Pándy-Szekeres et al[3].

Steric clashes were energy minimized using the MOE software[4]. Crystallographic waters and lipid modifications present in the X-ray structure were preserved. The rest of co-crystallized molecules were discarded. Receptor residue protonation and tautomeric states were assigned using PROPKA[5,6] as implemented in PDB2PQR[7] at pH 7 and subsequently curated by members of the GPCRmd consortium. Residue D2x50 was either kept deprotonated or protonated for antagonist- or agonist-bound receptor complexes, respectively.

*Ligand parameterization*. Tripos Mol2 File Format files were taken from the PDB using the MOE software[4]. Protonation and tautomeric states at pH 7 were predicted and assigned using the Marvin Calculator Plugin[8]. The obtained Mol2 files were used to generate parameters by analogy using the ParamChem server 1.0.0 (https://cgenff.umaryland.edu) and CGenFF 3.0.1[9–12]. Parameters for inorganic phosphate ($PO_4^{2-}$) were obtained from CGenFF and SwissParam[13] web server. Finally, adenosine parameters where taken from the RNA CHARMM 36 force-field[14]

*Placement of additional internal waters*: For each structure we used HomolWat (http://lmc.uab.cat/homolwat/) to incorporate internal water molecules not determined in other structures of the same or parent receptors. To this end, we used water molecules with a circular variance[17] > 0.6 measured using vectors from the oxygen atom of a water molecule to the surrounding atoms up to 10 Å. The algorithm uses *blastp* (ncbi-blast v2.6.0+)[16] to generate an ordered list of receptors with determined structures that contain resolved water molecules and subsequently tries to incorporate into the model all water molecules that do not clash (> 2.4 Å) with receptor atoms or previously introduced water molecules, starting with 1) receptors with the largest sequence identity, 2) structures with the best resolution and 3) water molecules with the smallest B-factors. Water hydrogens are added using the PDB2PQR software[7]. Non-coincident waters molecules (distance > 2 Å) predicted by Dowser+ software[18] were also incorporated into the structure.

*System set-up*. Structures generated by the GPCRmd consortium were checked for protonation consistency. Acetylated and charged N terminus were used for incomplete and complete N terminus capping, respectively. Amidated and charged C terminus were used for incomplete and complete C terminus capping, respectively. Each GPCR model was aligned to its respective orientation taken from the Orientations of Proteins in Membranes database[19] using STAMP 4.4[20,21] . Receptors were then embedded into a POPC bilayer and solvated ensuring a 20 Å distance between protein periodic distances, considering also receptor diffusional rotation. Furthermore, the system was checked for lipids inserted into aromatic rings. Finally, systems were solvated with TIP3 water molecules and the ionic strength of the solution was adjusted to 0.15 M using NaCl ions. Parameters for the simulation were obtained from the CHARMM36m force field[22,23].

*Molecular dynamics (MD) simulations*. Systems were first energy minimized during 5000 step and then equilibrated at constant pressure (NPT, 1.01325 bar) using the Berendsen barostat[24] with a pressure relaxation time of 800 fs and a compressibility factor of $4.57x10^{-5}$ $bar^{-1}$ during 30 ns. In a first step, harmonic restraints of 1.0

kcal/(mol·Å$^2$) were set on protein backbone and water oxygen atoms during 10 ns. Then, restrains were progressively released in a ramp of -0.095 kcal/(mol·Å$^2$·ns) during 10 ns followed by a restraints-free equilibration step of 10 ns. Production simulations were performed at constant volume (NVT) in 3 replicates of 500 ns per system using ACEMD[25] and GPUGRID[26]. Time-step of 2 and 4 fs were used during the equilibration and production runs, respectively. Non-bonded interactions were cut-off at 9 Å. A smooth switching function for the cut-off was applied, starting at 7.5 Å. Long-distance electrostatic forces were calculated using the Particle Mesh Ewald algorithm[27] with a grid spacing of 1 Å. Bond lengths of hydrogen atoms were kept constrained using the RATTLE algorithm[28]. All simulations were carried out at a temperature of 310K using the Langevin thermostat[29] with damping constants γ of 1 ps$^{-1}$ and 0.1 ps$^{-1}$ for NPT and NVT simulations, respectively.

### Supplementary Note B: GPCRmd platform structure and tools

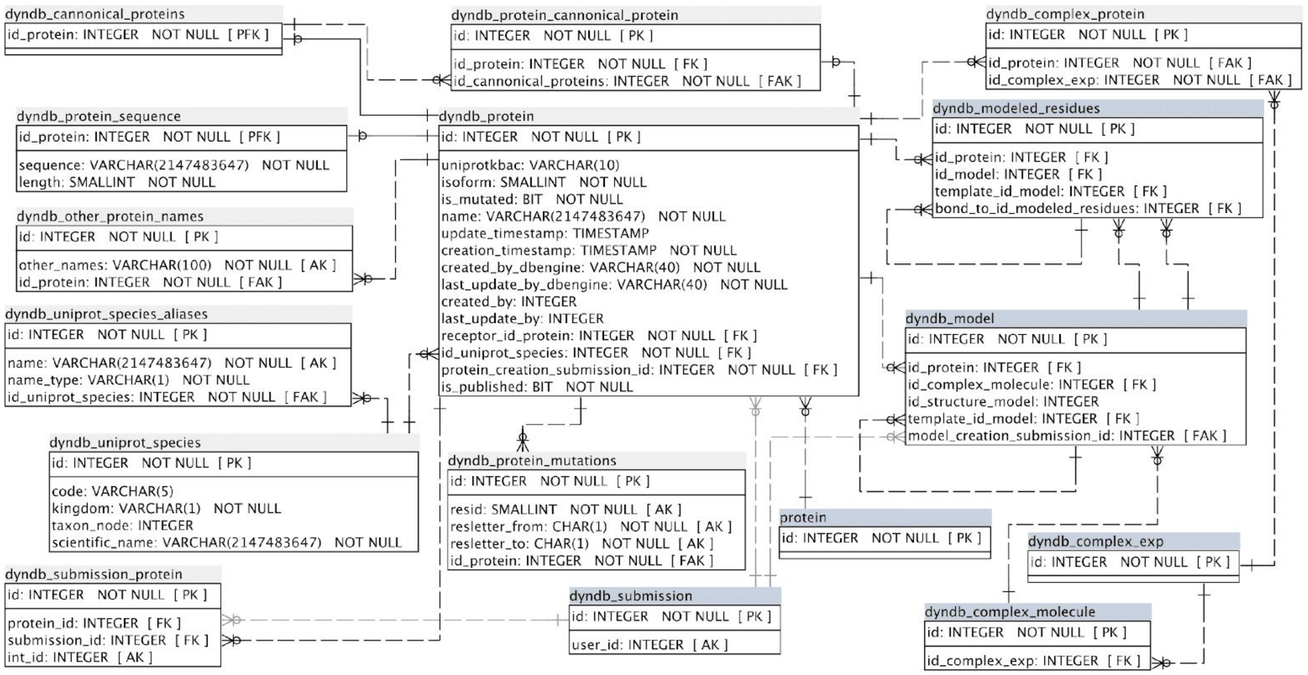

**Supplementary Figure 1**. GPCRmd entity-relationship (ER) diagram of entities related to protein objects. Tables in blue only display fields taking part in the relationships.

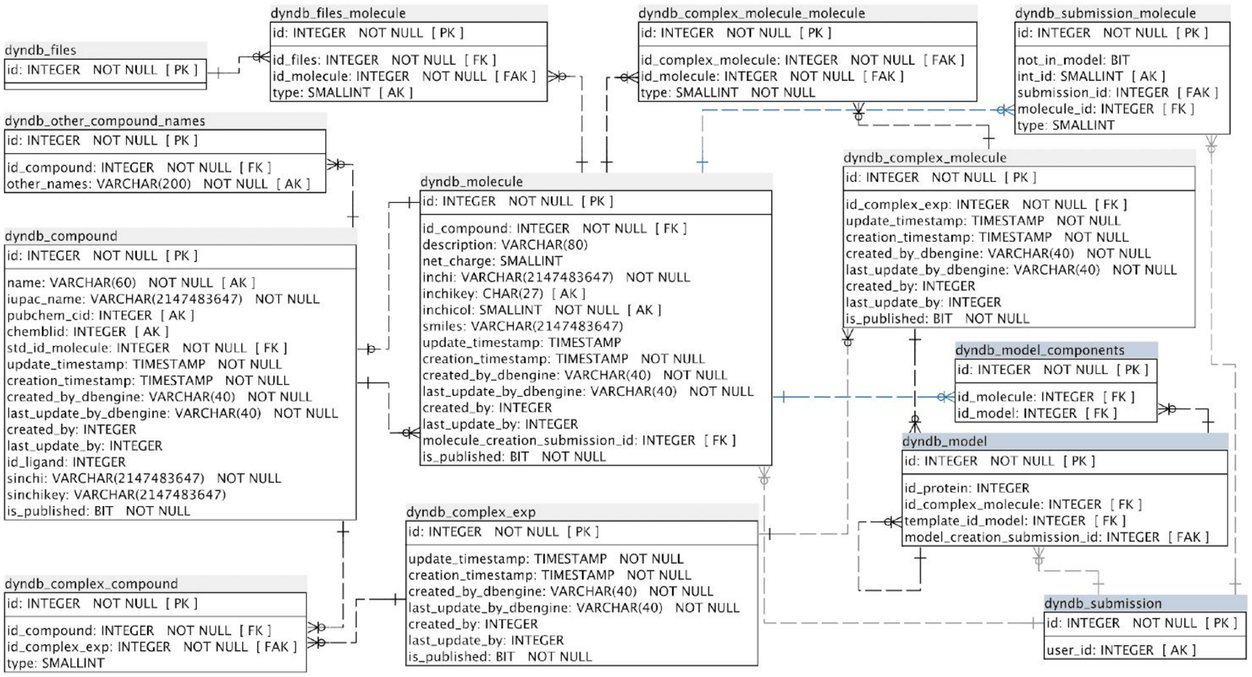


**Supplementary Figure 2**. GPCRmd ER diagram of entities related to molecule objects. Tables in blue only display fields taking part in the relationships.

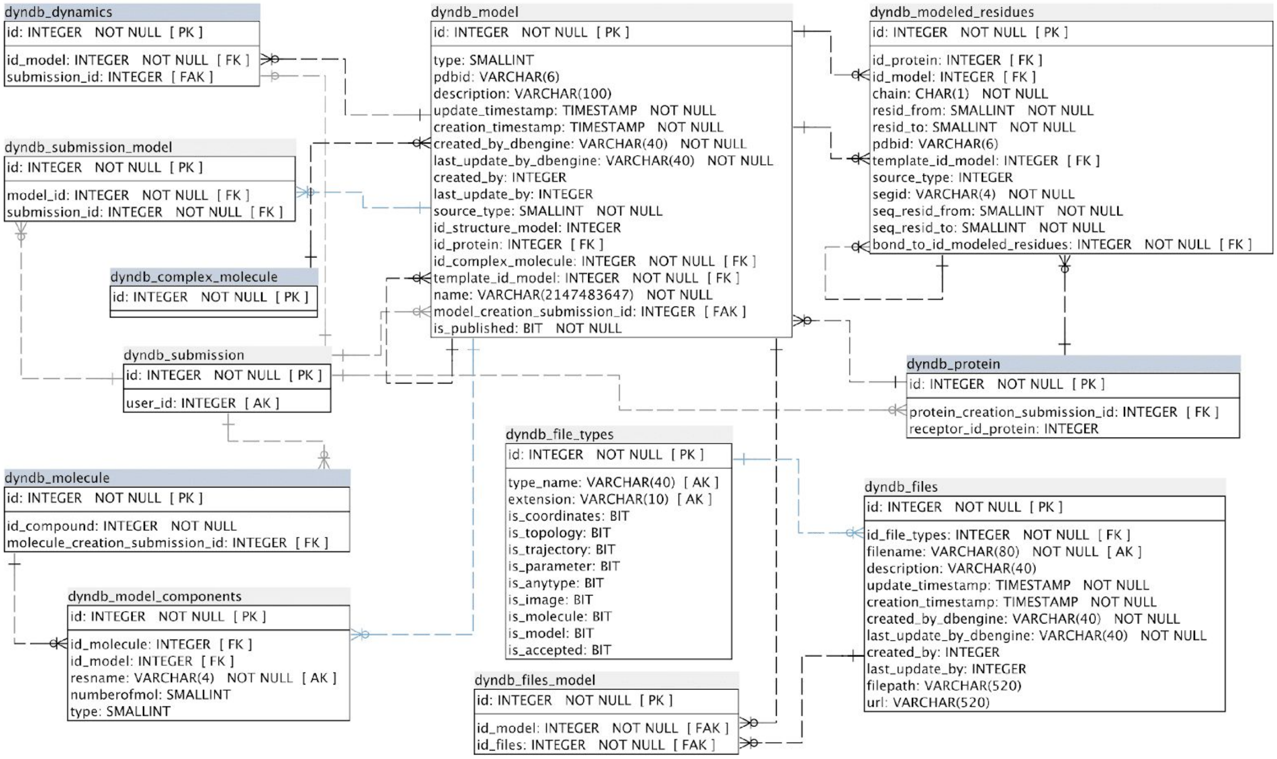


**Supplementary Figure 3**. GPCRmd ER diagram of entities related to model objects. Tables in blue only display fields taking part in the relationships.

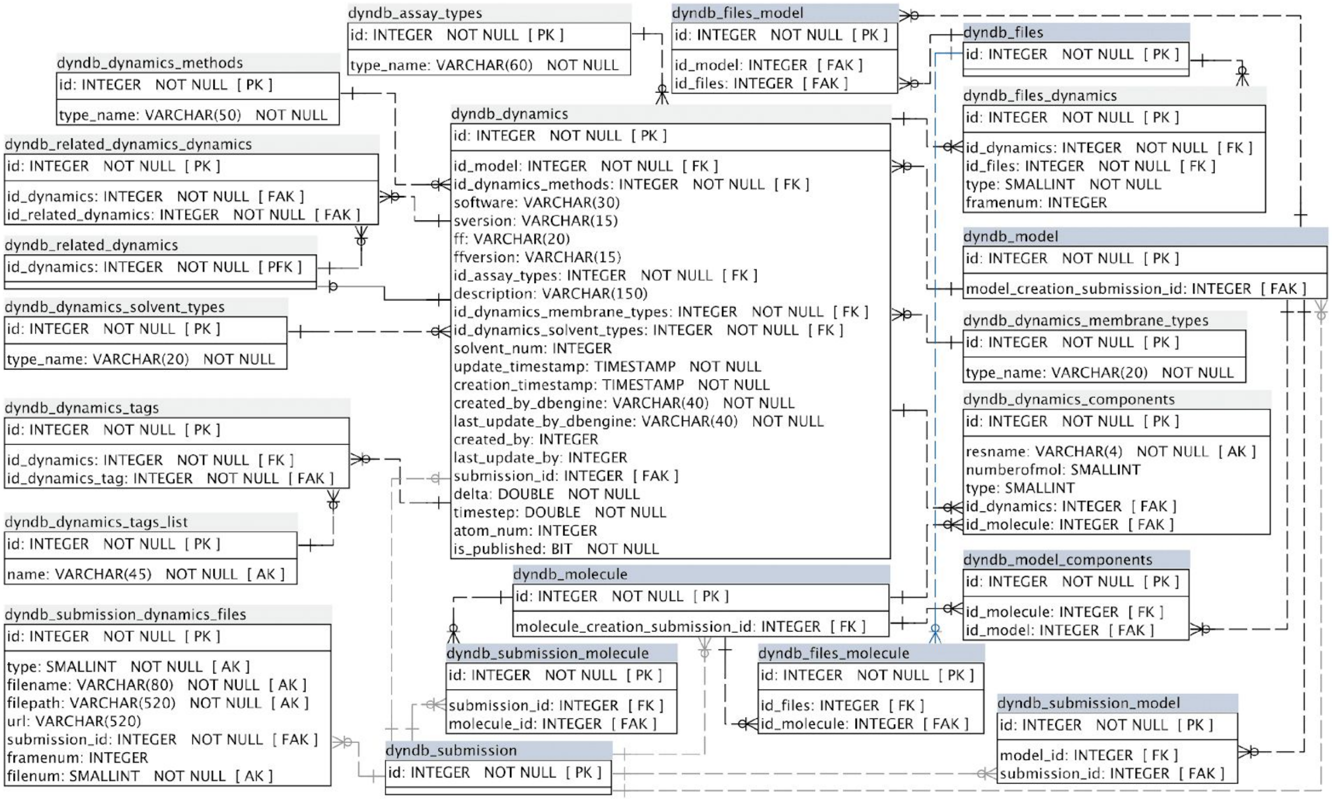


**Supplementary Figure 4**. GPCRmd ER diagram of entities related to dynamics objects. Tables in blue only display fields taking part in the relationships.

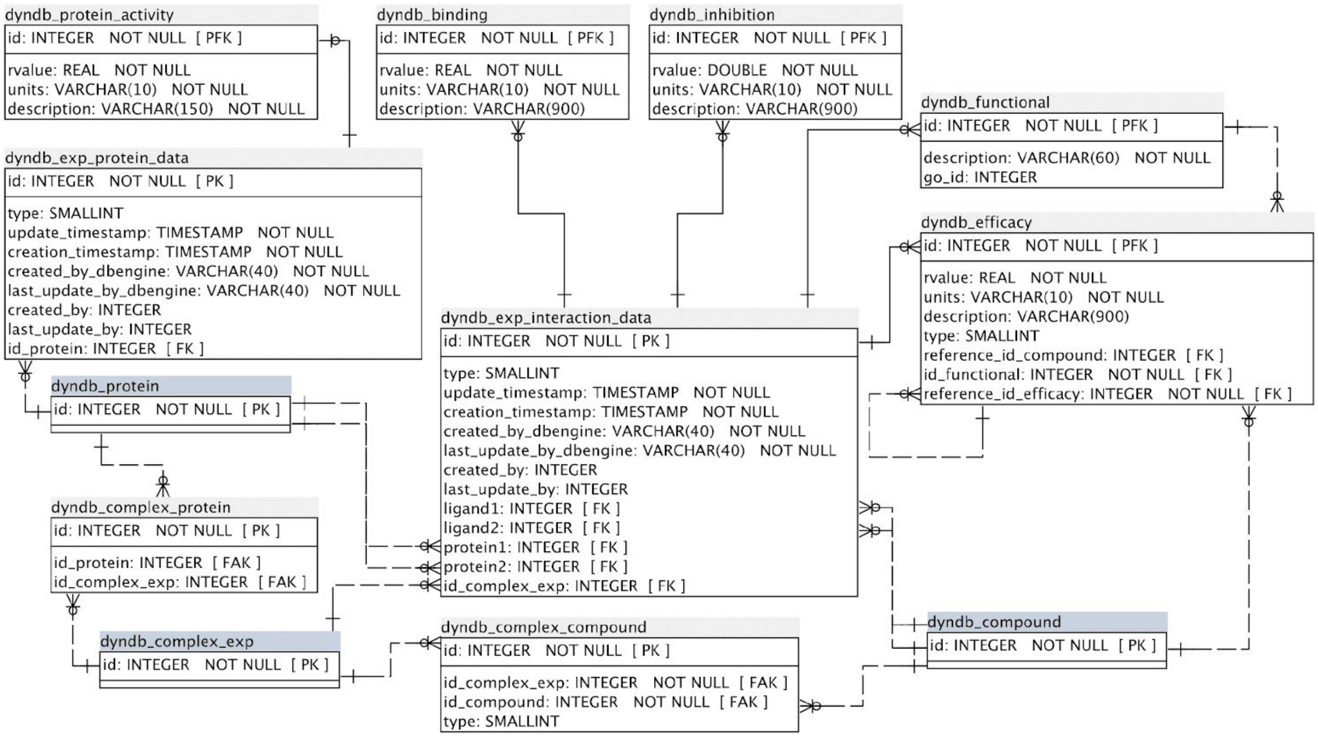


**Supplementary Figure 5**. GPCRmd ER diagram of entities related to experimental data. Tables in blue only display fields taking part in the relationships.

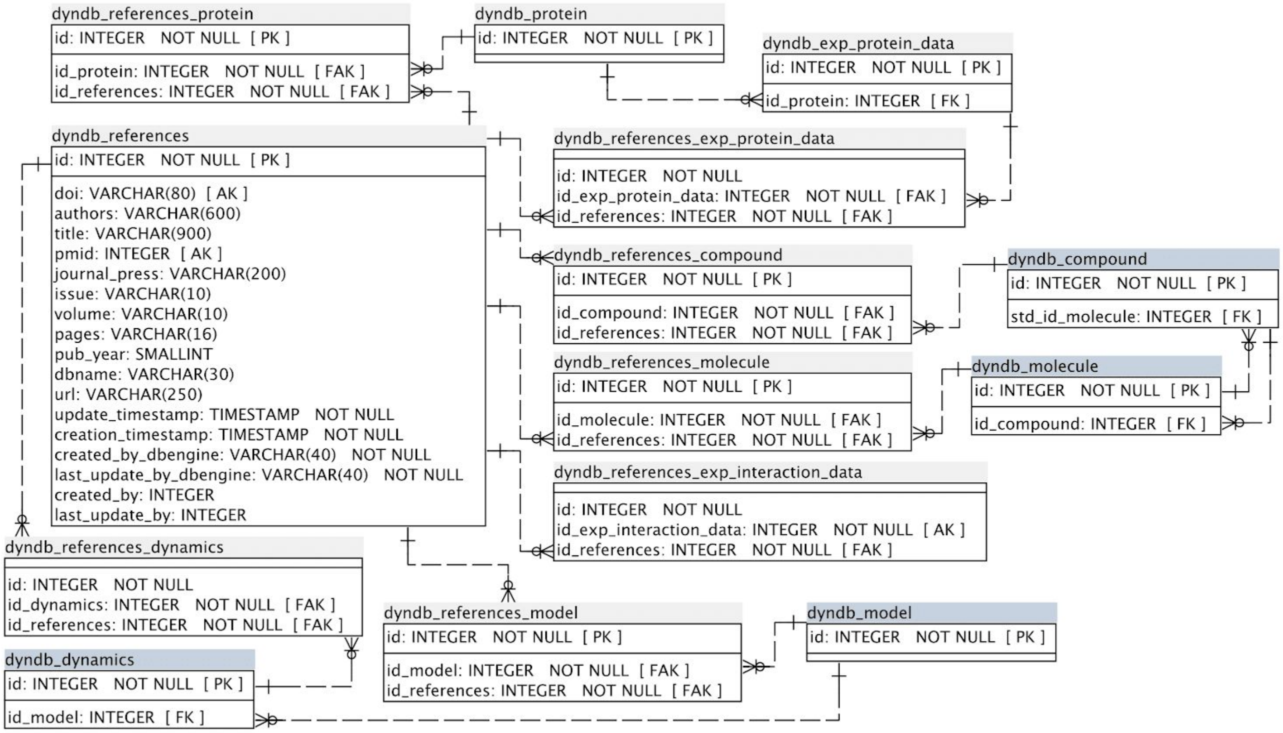


**Supplementary Figure 6**. GPCRmd ER diagram of entities related to references objects. Tables in blue only display fields taking part in the relationships.

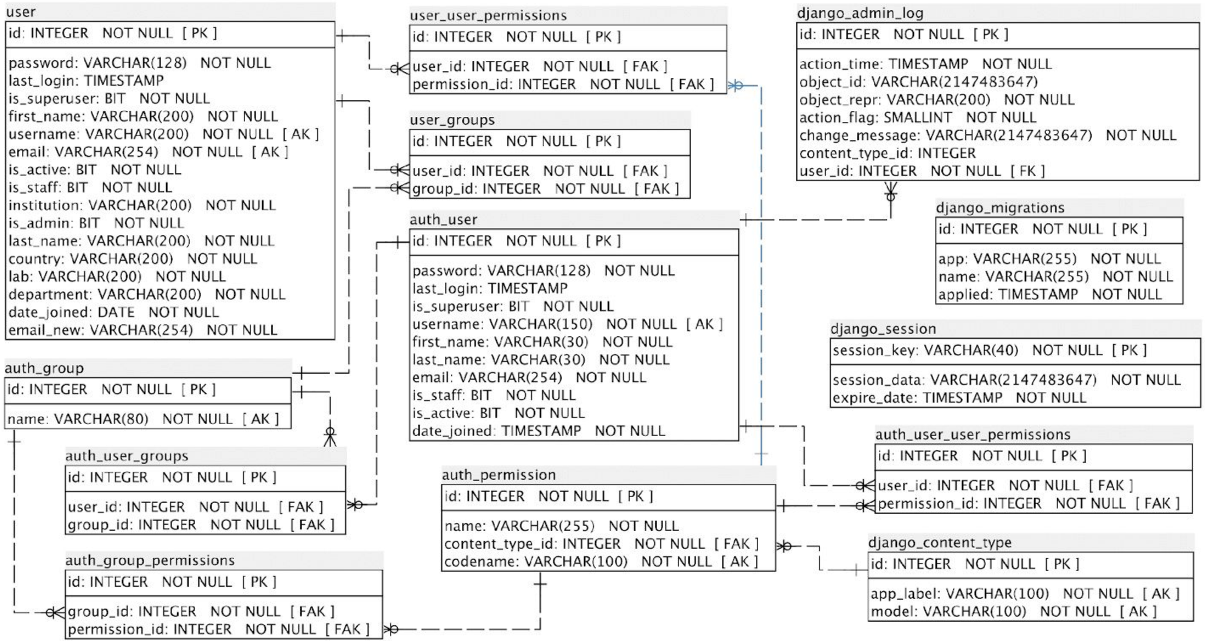


**Supplementary Figure 7**. GPCRmd ER diagram of user and Django tables.

**Supplementary Note C. Comparative analysis of water-mediated interaction**

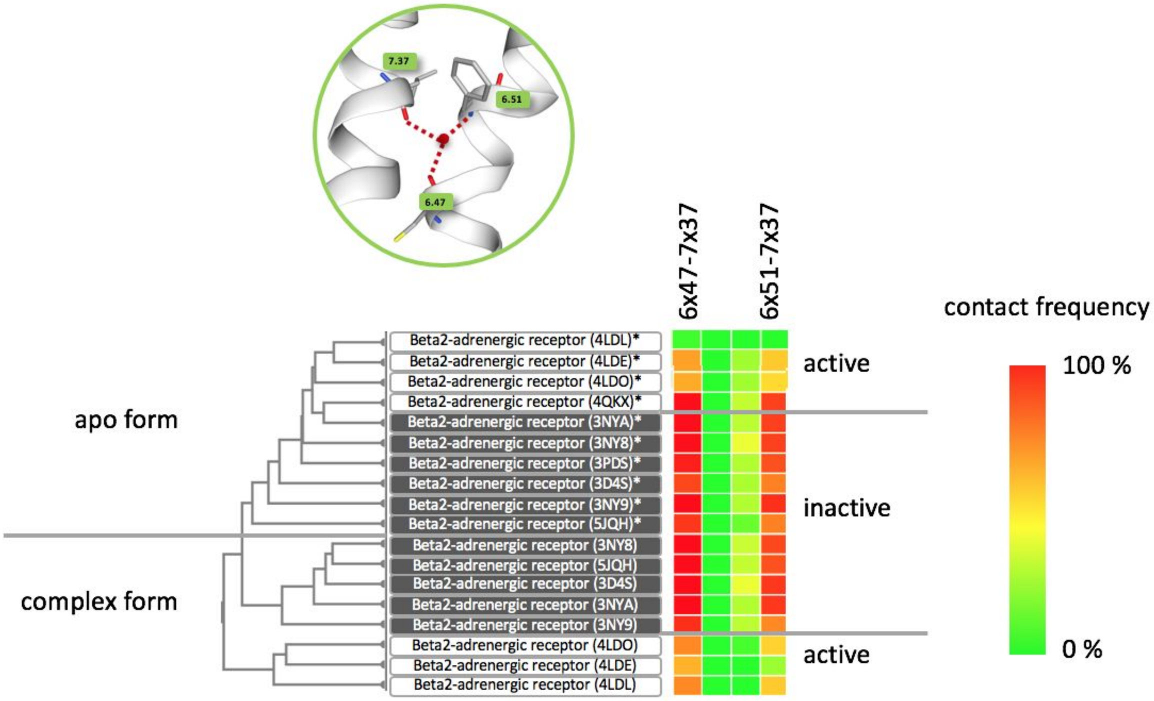


**Supplementary Figure 8**. Water-mediated contact map of the β2AR highlights the bifurcated network between TM6 and TM7. Systems marked with an asterisk indicate apo-form simulations which are nicely separated from the ligand-receptor complex simulations by the clustering analysis. Within the clustered groups for apo- and complex forms, we see also a separation of active and inactive structures. The bifurcated network which links TM6 to TM7 can be seen in inactive as well as active structures. Interestingly, contact frequencies are reduced in actives structures indicating a network loosening.

**Supplementary Note D. Sustainability**

Projects such as the GPCRmd face huge challenges to overcome in order to be accepted, used and sustained in the future. As previously shown, structural biology or (proteo-)genomics projects like the PDB (rcsb.org[1]) or the Galaxy project[30], respectively, have demonstrated the need for strong community support rather than individual laboratory to preserve an adequate sustainability. Therefore, we created the GPCRmd consortium under the umbrella of the newly granted ERNEST network (GLISTEN COST action, https://ernest-gpcr.eu), with the support of the GPCR community. Additionally, the focus on a specific research area such as GPCRs rather than a general database for all MD simulations reduces hurdles like deposition space problems, too general and likely unused analysis tools and problems in the findability due to too broad keywords/labels. An important asset promoting sustainability is the automated update of GPCRmd with new MD simulations for newly published GPCR structures. These updates will follow a standardized protocol developed by the GPCRmd community. On the other hand, GPCRmd is designed to serve as a revision platform in the near future allowing the editor and reviewers to evaluate dynamics data on the fly, making them more transparent and trustful and allowing upon acceptance for an easy deposition into the database.
